# Mitigation of intertidal temperature extremes by algal canopies

**DOI:** 10.64898/2026.10.29.762738

**Authors:** Cátia Monteiro, Carolina Miranda, Pedro Duarte-Coelho, Rita da Silva, Joana Pereira, Rocío Nieto-Vilela, Mar Humet, Ana Luísa Mano, Frederico Almada, Fernando P. Lima

## Abstract

In a rapidly warming world, species must either adapt to changing environmental conditions, shift their distributions through the movement and survival of individuals, or face local or global extinction. Within ecosystems, structural complexity of either environmental or biogenic origin may provide microclimate buffering, promoting persistence. In coastal environments, it has long been stated that macroalgae act as important ecosystem engineers that can moderate thermal extremes, however the magnitude of this buffering has rarely been quantified. Moreover, the extent to which thermal buffering varies among macroalgal species, and how it relates to thermal heterogeneity across scales remains poorly understood.

Here, we measured thermal buffering provided by intertidal macroalgae using high-frequency, high-spatial-resolution *in situ* temperature logging. We compared microclimates beneath algal cover with those on experimentally cleared bare rock across two tidal levels, two solar exposures and two seasons along six shores within a biogeographical transition zone.

Thermal buffering was highly heterogeneous across the study area, varying with species identity, algal cover, tidal level, and solar exposure. Together, these factors generated a complex thermal mosaic, with even small algal specimens providing measurable buffering.

We found that latitude had little influence on mean daytime low-tide temperatures, whereas local-scale factors had pronounced effects. Lower tidal position, reduced solar exposure, and algal cover each moderated thermal conditions and, in combination, provided an average cooling of approximately 4 °C. These findings highlight the importance of fine-scale habitat heterogeneity in shaping thermal exposure and identify features that could be incorporated into local climate-adaptation strategies, supporting coastal conservation and restoration efforts.

## Introduction

The rocky intertidal, lying between low and high tide, is a highly thermally complex ecosystem in which organisms must tolerate wide daily fluctuations in both air and water temperatures. The thermal landscape of the intertidal emerges from the interplay of processes operating across multiple spatial scales, ranging from large-scale latitudinal gradients and regional upwelling regimes to local factors such as geomorphology, solar exposure and biogenic habitats [1,2]. This complexity creates high variability in temperatures at the centimeter scale that is not captured by remote sensing. Hence, relying solely on satellite-derived sea surface temperatures to understand the thermal envelope of rocky shores has been shown to be insufficient [3–5]. *In situ* temperature monitoring is, therefore, key to understanding the ecological processes at play on rocky shores. Recent technological developments have overcome previous limitations in the durability, cost and precision of temperature sensors [6], making it possible to deploy a network of miniaturized temperature sensors *in situ* across the different microhabitats occurring on rocky shores, including crevices, beneath algae, and inside tidepools.

Because temperature is a major determinant of species’ biogeographical limits [7,8], this fine-scale thermal heterogeneity is also likely to influence patterns of species persistence and distribution. Gaps in our understanding of intertidal microclimates may therefore limit our ability to identify the mechanisms underlying ongoing biogeographical shifts.

Climate change-driven impacts are not occurring uniformly across space, creating valuable study areas where both highly diverse communities and thermal complexity occur. One of these study areas is the northwestern Iberian Peninsula, more specifically the Portuguese coastline. Over a coastline spanning little more than 600 km, intertidal communities shift from cold-adapted species in the north to temperate and warm-temperate assemblages in central and southern Portugal, respectively [9–11]. Large canopy-forming macroalgae such as *Ascophyllum nodosum*, *Fucus* spp. and *Bifurcaria bifurcata*, occur in the north, but they either disappear (*A. nodosum, B. bifurcata*) or become increasingly smaller and sparser towards the south (*Fucus* spp.)[12,13]. This biogeographical transition is driven not only by a latitudinal climatic gradient but, to a large extent, by the regional Canary upwelling system, which brings nutrient-rich cold waters to the surface in northern Portugal during summer [11,12]. Importantly, predictions of future upwelling scenarios include both decreased and increased upwelling intensity and duration [14,15], generating concerns over the persistence of these cold-adapted species in the near future. So far, range shifts have already occurred in some populations [16], whereas others appear comparatively stable [17]. Still, non-native species have increased and proliferated in the study area [18,19], with climate change frequently identified as an important underlying driver.

Importantly, biogenic habitats are common on rocky shores. Their role in facilitation, namely through thermal amelioration or buffering, is often stated, although with limited empirical evidence but see Rindi et al., (2026). Rocky shore biogenic habitats can consist of either animal aggregations (e.g., mussel beds or honeycomb reefs), or canopy-forming macroalgae. Canopy-forming macroalgae include kelps (large brown macroalgae of the orders Laminariales), that mostly inhabit subtidal environments, and fucoids and foliose red algae, in the intertidal. These canopy-forming macroalgae provide several ecosystem services and play important ecological roles, such as facilitation [21,22]. However, fucoids and foliose red algae have received far less attention than kelps [23], and hence, considerable knowledge gaps remain. Knowing how these intertidal algae contribute to the thermal landscape of their environment is key to understanding the temperatures actually experienced by organisms at their scale, which underpin ongoing and future ecosystem dynamics, and can dramatically improve species distribution models and projections of future distributions [24].

The objective of this work was to describe the thermal landscape of intertidal rocky shores by quantifying the thermal buffering provided by macroalgae across different spatial scales (latitude, tidal level, solar exposure) and temporal scales (season). We set up a comprehensive temperature sensor network targeting different macroalgae species and morphologies common along the Portuguese coastline, representative of a typical cold-to-warm temperate transition in macroalgae communities worldwide [25,26].

## Material and Methods

To assess the effect of macroalgal cover on intertidal temperature and thus its potential to provide thermal refugia for intertidal species, we deployed autonomous *in situ* temperature loggers (EnvLoggers T7.3, electricblue.eu), recording hourly data with a precision of 0.1 °C (Pereira et al., 2026). Temperature loggers were deployed in two treatments – “algal-cover” and “bare-rock”. In the “algal-cover” treatment, macroalgae were kept intact, and the loggers were deployed under their fronds. In the “bare-rock” treatment, macroalgae were removed, the substrate was scraped in an area of 25 x 25 cm, and loggers were deployed in the center of the plot, in bare rock. On shores where the algae form canopies, “algal-cover” loggers were deployed directly below the canopy next to the holdfast and were thus immediately covered by algae. In plots with algal turfs, hole drilling destroyed the natural algal cover, and the logger remained uncovered until the algae grew back. The two treatment levels were always set up in pairs to ensure that all other factors (e.g., tidal level, aspect, and slope) remained the same. In all cases, loggers were deployed into 12-mm deep, 30-mm diameter holes drilled into the rock, flush with the rock surface (Fig. S1).

The experiment was set up at six shores along the Portuguese coast between September 2022 and March 2024 (Fig. 1, Table S1). At each shore, three replicate pairs of “algal-cover”/”bare-rock” plots were set up at two tidal levels (top and low intertidal) and under two levels of solar exposure. “Hot” plots were located on south-oriented rock faces receiving maximum direct sunlight, whereas “Cold” plots were located on north-oriented or shaded surfaces receiving minimal direct sunlight. This design resulted in 24 plots per shore and 144 plots overall.

**Figure 1.**
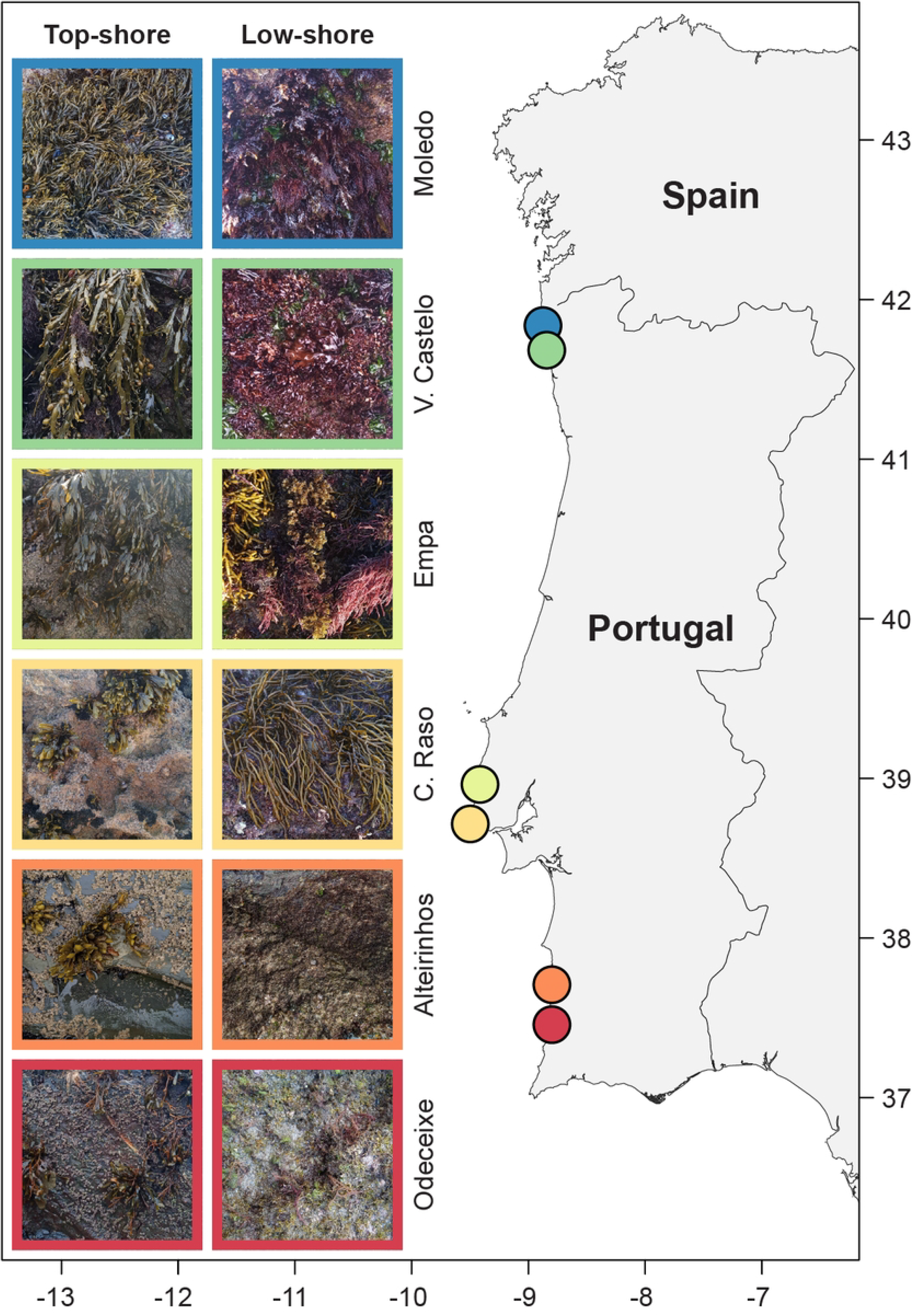
Map showing the location of the six shores surveyed in this study. Examples of the macroalgae targeted at the top and low intertidal levels at each shore are shown on the left. See Table 1 for more details.

**Table 1.**
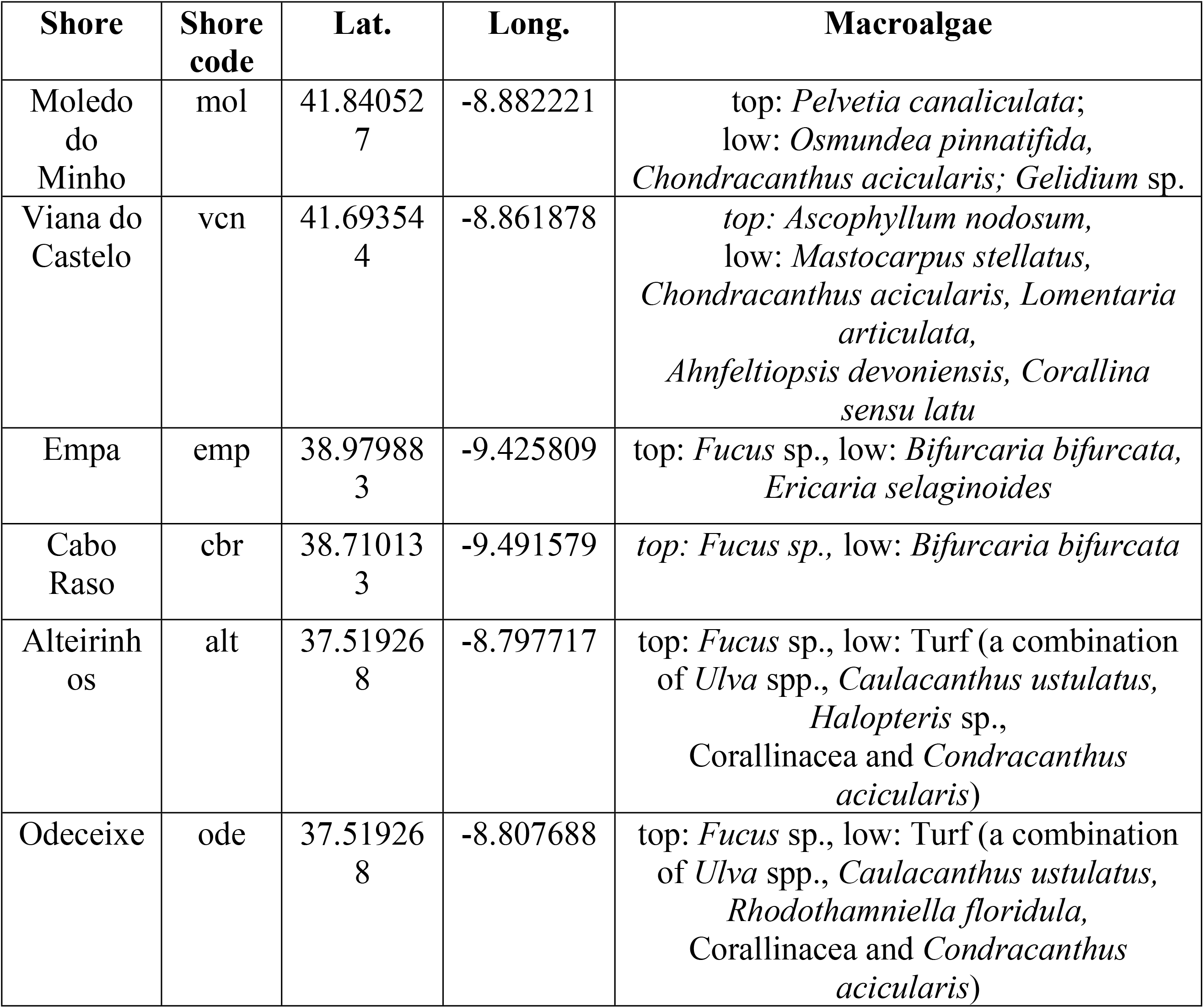
Shores sampled, the respective shore code and coordinates, and the seaweed taxa targeted in each shore and tidal level.

| Shore | Shore code | Lat. | Long. | Macroalgae |
|---|---|---|---|---|
| Moledo do Minho | mol | 41.840527 | -8.882221 | top: *Pelvetia canaliculata*;<br>low: *Osmundea pinnatifida,*<br>*Chondracanthus acicularis; Gelidium* sp. |
| Viana do Castelo | vcn | 41.693544 | -8.861878 | *top: Ascophyllum nodosum,*<br>low: *Mastocarpus stellatus,*<br>*Chondracanthus acicularis, Lomentaria articulata,*<br>*Ahnfeltiopsis devoniensis, Corallina sensu latu* |
| Empa | emp | 38.979883 | -9.425809 | top: *Fucus* sp., low: *Bifurcaria bifurcata, Ericaria selaginoides* |
| Cabo Raso | cbr | 38.710133 | -9.491579 | *top: Fucus sp.,* low: *Bifurcaria bifurcata* |
| Alteirinhos | alt | 37.519268 | -8.797717 | top: *Fucus* sp., low: Turf (a combination of *Ulva* spp., *Caulacanthus ustulatus, Halopteris* sp.,<br>Corallinacea and *Condracanthus acicularis*) |
| Odeceixe | ode | 37.519268 | -8.807688 | top: *Fucus* sp., low: Turf (a combination of *Ulva* spp., *Caulacanthus ustulatus, Rhodothamniella floridula,*<br>Corallinacea and *Condracanthus acicularis*) |

The identity of the algae covering the temperature loggers changed among locations, reflecting the natural variability along the study area (Fig. 1, Table 1). At Viana do Castelo and Moledo in northern Portugal, the brown canopy-forming macroalgae *Ascophyllum nodosum* and *Pelvetia canaliculata* were targeted in the top intertidal, while red foliose algae were targeted in the low intertidal. In central and southern Portugal, the brown macroalgae *Fucus* spp. were common in the top intertidal. In the low intertidal, medium-sized macroalgae such as *Bifurcaria bifurcata* and *Ericaria selaginoides* were common at Empa and Cabo Raso whereas only turf algae were common at Odeceixe and Alteirinhos.

Shores were visited approximately every two months. On each visit, pictures of all plots were taken, temperature data were downloaded, and loggers located on bare rock were scrubbed clean. Logger cover was assessed from photographs taken during each visit using the following ordinal scale: 1, clean; 2, slightly covered; 3, partially covered; 4, covered; 5, fully covered; 6, covered by limpets; 7, covered by *Sabellaria alveolata*.; 8, covered by sand; 9, covered by cobbles (Fig. S2). Based on field observations during the experiment, bare-rock loggers were assumed to remain clean for four weeks after each maintenance visit. Only algal-cover/bare-rock pairs for which algal cover on the algal-cover logger was scored as above 3 were retained for downstream subsequent analysis.

### Statistics

The number of logger pairs with algal cover scores above 3 (partially covered) was highly heterogeneous across shores and treatments, limiting the duration of the time series available for direct comparison. We therefore selected two discrete periods during the warmer part of the year, chosen to maximise the number of replicate pairs with adequate algal cover across shores and experimental conditions (Table S1). These periods occurred between 15 or 16 September 2023 and 13 or 14 October 2023 and are hereafter referred to as “summer”. Although September and October do not correspond strictly to the climatological summer period, high-temperature peaks and warm spells frequently occur during these months. As a reference, concerning average temperatures only, in 2023, the mean maximum air temperature in continental Portugal was 29.5 °C in August, 26.5 °C in September and 24.5 °C in October [27]. For winter, the selected periods varied among shores to maximise the number of algae-covered pairs (Fig. S2, Table S2). For each selected period, hourly logger data were classified by tidal state (low tide/high tide) and daylight (day/night) using predicted tide height and sunrise/sunset times. Height relative to mean sea level (MSL) was measured for the experimental plots and loggers using a centimetre-precision Global Navigation Satellite Systems (GNSS) receiver (SparkFun RTK Facet L-Band, Sparkfun, USA) with Real-Time Kinematic (RTK) corrections obtained from the nearest NTRIP-broadcasting station of the ReNEP network (Direção-Geral do Território, Portugal), and was used to classify high tide (water) and low tide (air) temperature records. Sunrise/sunset times were computed using the suncalc R package[28] from the geographic coordinates of each shore and were used to classify each logger timestamp as daytime or nighttime.

The buffer provided by algal cover was quantified for each pair as the temperature measured in the algae-covered plot minus the temperature in the bare-rock plot. Negative values therefore indicate cooling beneath the algae and positive values indicate warming. Solar exposure buffering was quantified as the temperature difference between cold microhabitats (north-facing or shaded surfaces) and hot microhabitats (south-facing, sun exposed surfaces), using bare-rock plots only. Negative values indicate that cold microhabitats were cooler than hot microhabitats.

To reduce temporal pseudoreplication, hourly daytime low-tide temperatures were aggregated into daily means for the main analysis and into daily 90^th^ percentiles for the analysis of temperature extremes. Mixed-effects models [29] were fitted to test the effects of season, tidal level (top shore vs low shore), solar exposure (hot/cold), latitude (six locations from 41.84° to 37.52°), bare-rock temperature, and buffer type, with repeated structure by shore/date where possible. Latitude effects were also evaluated using shore-level summaries and interpreted cautiously because only six locations were sampled. Finally, model predictions were used to partition temperature reductions from a hot, top-shore, bare-rock baseline into season-specific contributions of tidal level, solar exposure, and algal cover. The analyses were run in R version 4.6.1.

## Results

### Seasonality of macroalgal cover and methodological constraints

During the study period (between September 2022 and March 2024), macroalgal cover varied at each shore (Fig. S2). During the spring, algal cover increased at almost all shores, reaching a maximum by the end of summer, after which some patches of macroalgae disappeared. On several shores, algae were replaced during winter by *Sabellaria alveolata* reefs. *Ulva* sp. was also quick to occupy space released by algae removal in bare-rock plots during autumn. The only exception was in Viana do Castelo, in northern Portugal, where *Ascophyllum nodosum* cover remained relatively stable through the year, albeit the extent of cover varied somewhat over time. In central Portugal (Cabo Raso and Empa), the patchy distribution and small size of *Fucus* spp. Canopies (consisting of only a small number of individuals a few cm in size) made it difficult to maintain a continuous cover over the temperature loggers. At these shores, *Fucus* spp. cover disappeared in January 2023 and did not recover during the remainder of the experiment. A similar pattern was observed for *Pelvetia canaliculata* in Moledo do Minho, in northern Portugal, where all temperature loggers became uncovered from October 2023 onwards. In several plots, *P. canaliculata* recruited in the immediate vicinity of the loggers but did not subsequently cover them.

### Effect of latitude on intertidal temperatures

We found no significant effect of latitude on intertidal temperatures (95% CI: -0.453 to 0.097, p=0.147, Fig. S3). This result, however, should be interpreted with caution as only six shores were studied, spanning less than 4 degrees in latitude.

### Effect of local scale factors on intertidal temperatures

Both algal cover and solar exposure modulated temperature during diurnal low tides (red dots, Fig. 2), despite having a much smaller effect on temperature during high tide and during nocturnal low tides (blue dots, Fig. 2), both during summer and winter (top and bottom panels, respectively). The thermal buffering offered by algal cover was higher in hot microhabitats than in the cold microhabitats.

**Figure 2.**
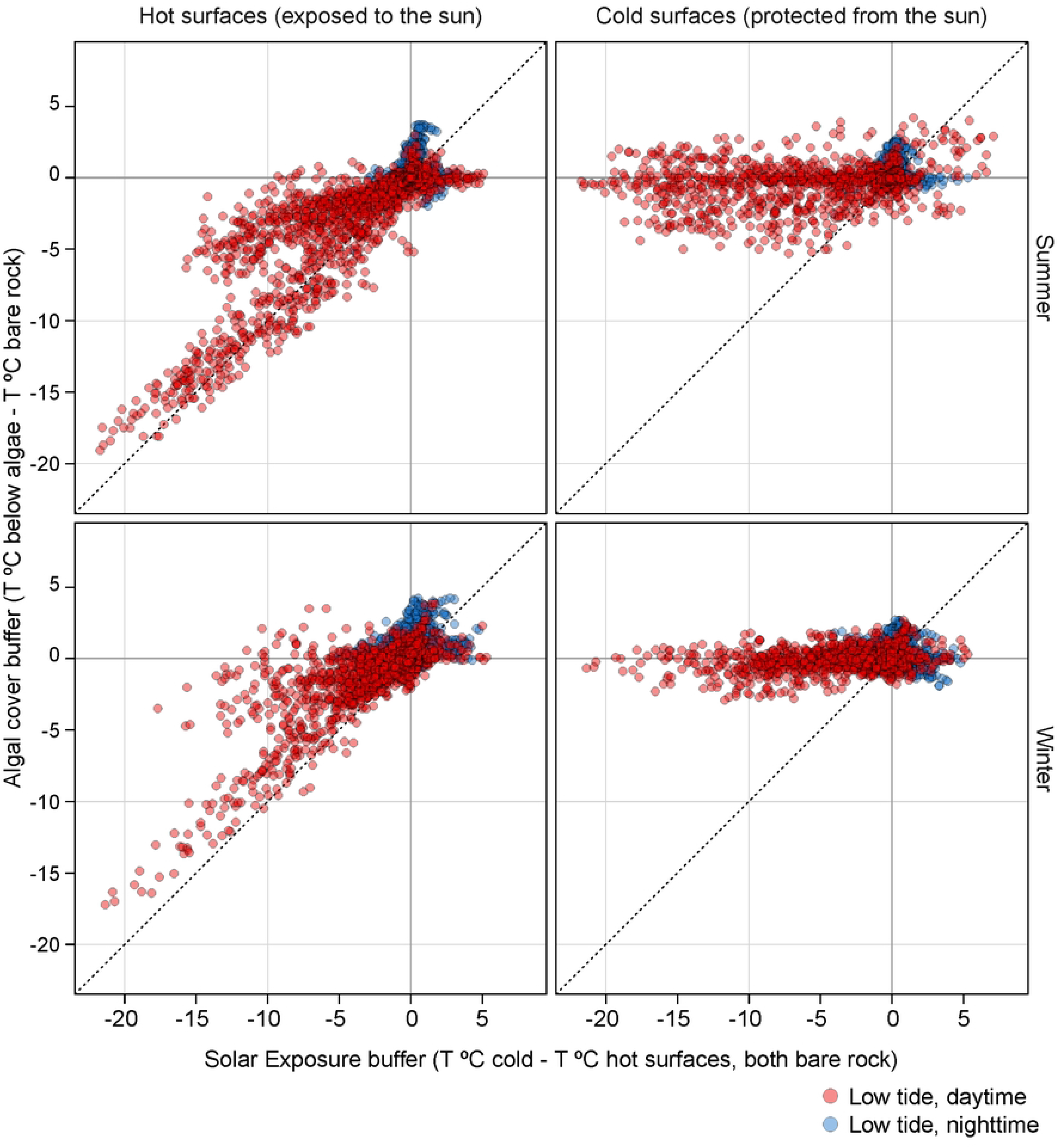
Scatter plots comparing algal cover buffer (temperature beneath algae minus temperature on bare rock) with solar exposure buffering (temperature in cold minus hot bare-rock microhabitats) in summer (top) and winter (bottom), and in hot (left) and cold (right) microhabitats. Data from both tidal levels are presented together. Red dots and blue dots represent daily mean low-tide temperatures during the day and during the night, respectively.

Mixed-effects models revealed that the local scale factors tidal level, solar exposure, and algal cover had significant effects on mean daytime low tide (during emersion) temperatures. The effect of tidal level was the strongest – temperatures at low level were, in average, 1.31 °C lower than in the top (95% CI: -1.46 to -1.15, p < 0.001); followed by solar exposure – temperatures at cold microhabitats where, on average, 0.8 °C lower than hot microhabitats (95% CI: -0.825 to -0.778, p < 0.001); and algal cover – temperatures beneath algae were, on average, 0.52 °C lower than on bare rock (95% CI: -0.532 to -0.497, p < 0.001). The coolest microclimate conditions occurred where these factors coincided: low shore, cold microhabitat, and algal cover.

Thermal buffering by solar protection and algal cover was stronger for extreme temperatures, represented by the daily 90^th^ percentile, than for daily mean temperatures. For the 90^th^ percentile, the effect of solar exposure was -0.83°C (95% CI: -0.850 to -0.813, p < 0.001), whereas the effect of algal cover was -0.53°C (95% CI: -0.549 to -0.513, p < 0.001).

Solar exposure (cold minus hot microhabitats) had a stronger buffering effect than algal cover (algal cover minus bare rock) (Fig. 3). For daily mean daytime low-tide temperatures, solar exposure buffering was on average 0.47 °C greater than algal-cover buffering (95% CI: 0.418 to 0.522, p< 0.001). Under extreme hot conditions, represented by the 90^th^ percentile, this difference increased to 0.84 °C (95% CI: 0.748 to 0.926, p< 0.001). However, the relative contribution of algal cover and solar exposure varied among shores.

**Figure 3.**
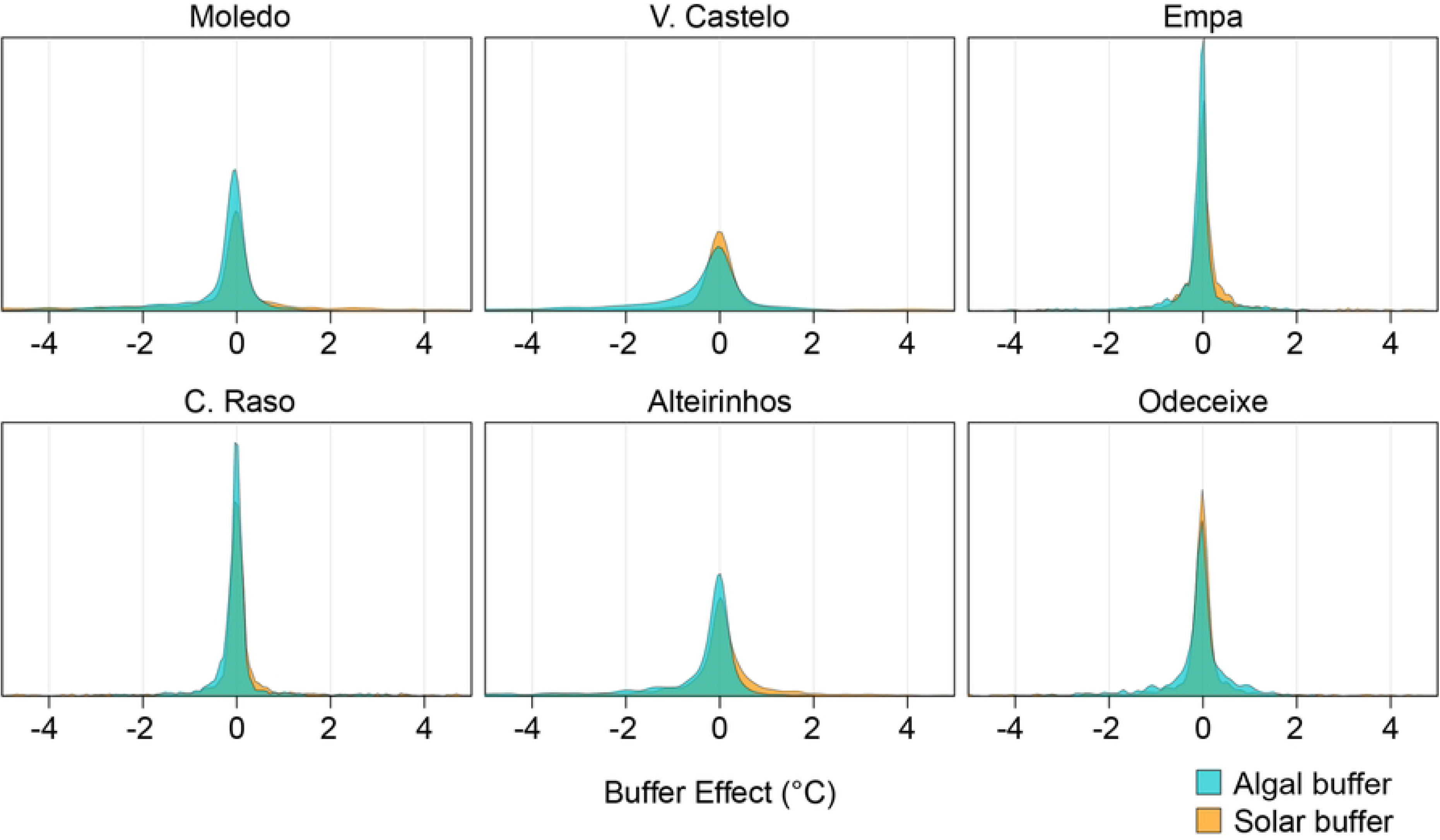
Ridge plots comparing the buffering effect of algal cover (yellow) and solar exposure (blue) at every shore, ordered from north (left top) to south (right bottom). Data from both tidal levels are presented together. Algal-cover buffering was calculated as temperature beneath algae minus temperature on bare rock, and solar exposure buffering as temperature in cold minus hot microhabitats.

Figure 4 shows the combined thermal effects of algal cover buffer, solar exposure, and tidal level at Viana do Castelo. Regression estimates are provided in table S3. Equivalent plots for all shores are shown in Fig. S4. At Viana do Castelo, bare-rock temperatures exceeded 40 °C in top-intertidal hot microhabitats. At these bare-rock temperatures, temperatures beneath algal cover were up to 20 °C lower, representing the strongest algae buffering recorded in the study. Algae buffering was highest in the top-intertidal hot microhabitats and lowest in low-intertidal cold microhabitats. The same overall pattern was observed at the remaining shores, although the magnitude of buffering differed (Fig. S4). At these shores, maximum bare-rock temperatures also reached approximately 35 °C to 40 °C, but the maximum algae-buffering effect was approximately 5 °C (Fig. S4).

**Figure 4.**
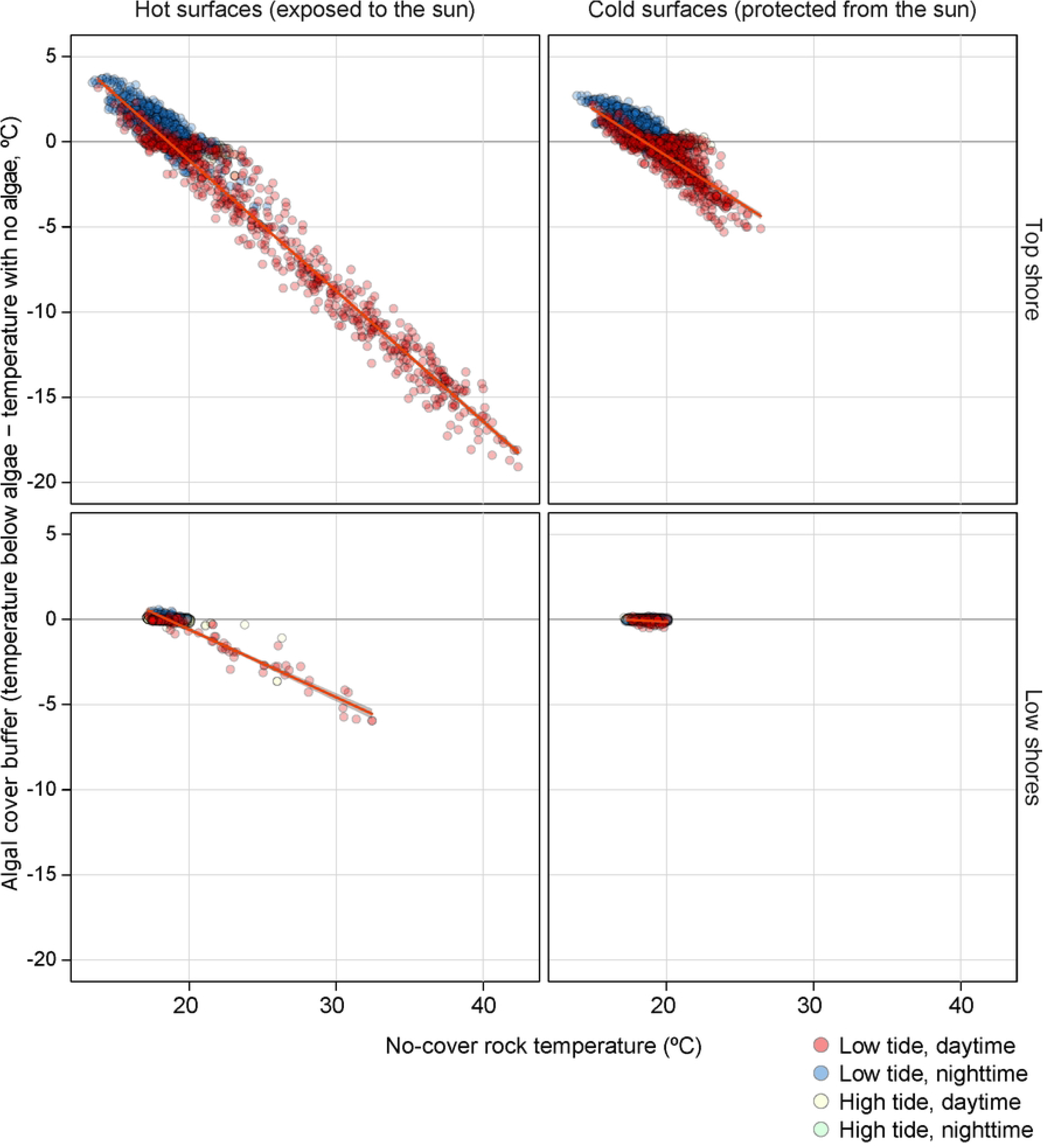
Relationships between algal-cover effect and bare-rock temperatures at Viana do Castelo (vcn). Panels from left to right show the tidal level x solar exposure combinations: top hot, top cold, low hot, low cold. Daily mean temperatures are represented by red dots (low tide, daytime), blue dots (low tide, nighttime), light yellow dots (high tide, daytime) and light green dots (high tide, nighttime).

The partitioning analysis estimated the combined effect of the local-scale factors on realized microhabitat temperatures. Moving from the hottest combination of conditions (top intertidal, hot exposure, and bare-rock) to the coolest combination (low intertidal, cold exposure, beneath algae) resulted in a predicted reduction of 3.99 °C in daily mean daytime low-tide temperature, and 8.46 °C in the daily 90^th^ percentile (Fig. 5).

**Figure 5.**
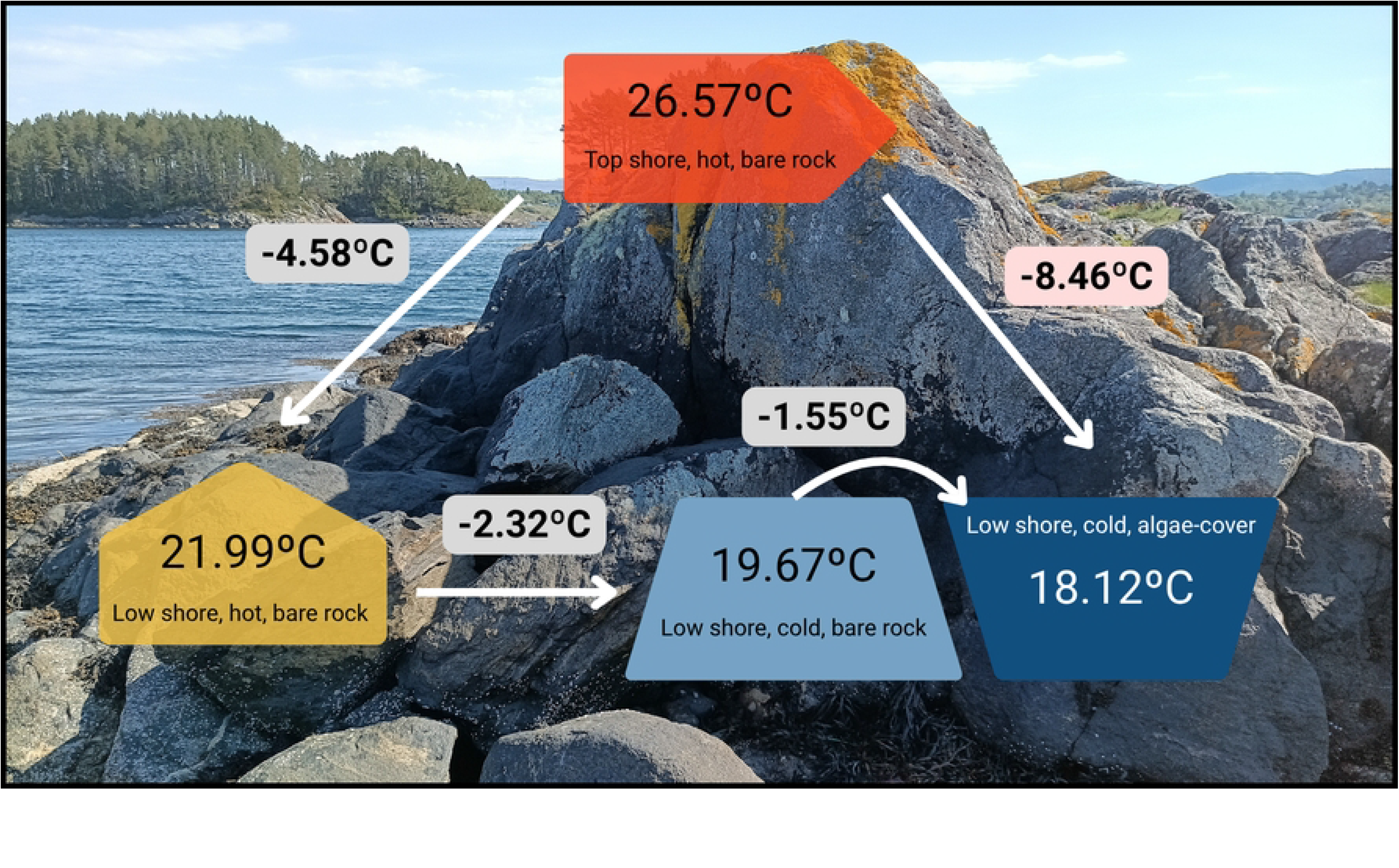
Schematic representation of the partitioning analysis. The predicted summer daytime low-tide 90^th^ percentile temperature for a hot, top-intertidal, bare rock microhabitat at a mean latitude of 39.67 °N was 26.57 °C. Moving to low intertidal reduced this temperature by 4.58 °C, moving to a cold microhabitat reduced it by a further 2.32 °C, and adding algal cover reduced it by an additional 1.55 °C. The cumulative predicted reduction from a hot, top-intertidal, bare-rock microhabitat to a cold, low-intertidal, algal-covered microhabitat was therefore 8.46 °C.

## Discussion

This study quantifies the thermal buffering offered by algal cover in comparison with bare rock (scraped clean regularly) in the intertidal across different local-scale conditions, including tidal level and solar exposure, and across a biogeographical transition zone in the North Atlantic, encompassing six shores in northern, central and southern Portugal. Hence, we provide unprecedented spatial resolution for measurements of intertidal temperatures below algal cover [30]. Algal cover reduced temperatures under most conditions tested. Thermal buffering was highest in summer, at the top intertidal and in hot microhabitats, which correspond to the most thermally stressful conditions, highlighting the potential of macroalgae to act as local climate refugia. Nevertheless, algal buffering persisted even in less extreme environments (e.g., the low intertidal, in cold microhabitats, and during winter) and even when canopies were small and sparse (e.g., in southern Portugal).

Although the role of canopies as a thermal buffer is well established in terrestrial systems [31,32], few studies have consistently measured this effect in coastal environments [22]. Moreover, in coastal environments, this research has focused on large brown canopy-forming macroalgae [20,30], while other macroalgae, such as red foliose algae or smaller brown algae, have been largely ignored (but see [13,33]). By applying a standardized approach to several Rhodophyta and Phaeophyta taxa, our study shows that thermal buffering is not restricted to large macroalgal canopies. The taxa selected at each shore also reflected the natural biogeographical transition in macroalgal communities along the Portuguese coast [9].

Western Iberia is positioned within the Canary upwelling system, which strongly influences coastal temperatures during summer [34]. The cold and nutrient-rich water brought to the surface from depth by summer upwelling creates a stronghold of cold-affinity species in the northwestern Iberia (northern Portugal and Galicia) [10] that differs strongly from the communities of central and southern Portugal, which are mainly populated by warm-water species [11]. This biogeographic transition is captured in this study by the larger specimens and more abundant canopies of brown algae studied at the top intertidal level in northern, at Moledo and Viana do Castelo (*Pelvetia canaliculata* and *Ascophyllum nodosum*), compared to smaller and less voluminous canopies of *Fucus* sp. present at shores in central and southern Portugal. In the low intertidal level, species targeted ranged from large foliose red algae in northern Portugal to turf algae in southern Portugal.

Importantly, all targeted species provided some degree of thermal buffering when algal cover was present. The thermal buffering provided by algae in the southern region highlights their potential to provide less stressful environments for species that are already near the edge of their distribution ranges. At the southern shores, macroalgae are not only shorter (i.e., with only a few centimeters in height) but are also sparser, providing limited algal cover. Still, a cooling of up to 7.5 °C was measured immediately below *Fucus* spp. in the top and up to 8 °C below turf algae (including *Rhodothamniella floridula* and *Ulva* sp., among others[35] in the low intertidal. This is consistent with a previous study, from the south of Portugal, which reported that temperatures beneath *Fucus* sp. in the upper tidal level were, on average, 9 °C lower than those recorded on the bare rock substrate [13]. This thermal buffering is probably associated with the spatial heterogeneity of macroalgae patches at the microscale (millimeters to centimeters) [36] which has been shown either to facilitate the presence of macroinvertebrate species [37] or to provide habitat comparable to that of larger canopy-forming algae for fish communities [38]. As large canopy-forming brown macroalgae are likely to retreat under climate change [39,40] and decrease in size [41,42], the microhabitat created by turfs and foliose algae needs to be better assessed, especially in tropical, warm-temperate environments where turfs are common, to improve our understanding of ecological and biogeographical patterns.

High seasonal variability in canopy extent was observed, except for the fucoid *Ascophyllum nodosum*. Canopies of *A. nodosum* were present throughout the year as expected, since *A. nodosum* is a long-lived, dominant intertidal species in the North Atlantic [43]. For other species, the seasonality observed here was surprising, as is the case for *Pelvetia canaliculata*. Although this species is considered perennial [44], in this study it displayed high variability in its cover at small (cm) scales, growing, disappearing and re-growing in patchy patterns around and on top of the experimental plots throughout the whole year. This was also very common for *Fucus* sp. in central and southern Portugal, where loggers assigned to the algal-cover treatment spent several months uncovered. These observations give important ecological insights into the role of macroalgae as climate refugia and habitat providers, as species will only benefit from these ecological roles if they can cope with the temporal variability in macroalgae presence and cover and/or if algal cover is present during particularly stressful events (e.g. heatwaves). Hence, the phenology of algal cover and its effects on the survival and performance of associated species need to be further explored. Importantly, given that sea surface and air temperatures along the Portuguese coast have risen and are expected to rise further with climate change [45–47], the buffer provided by these macroalgae may become increasingly relevant. Hence, including *in situ* thermal buffer measurements such as those obtained in this study in predictions of future species distributions could provide a more accurate picture of future communities under climate change and support the selection of sites and species for conservation and mitigation actions [2,48].

An important implication of these findings is that microclimatic buffering may complicate how climate-change-driven range shifts are detected and interpreted. Species occurring near the warm edge of their distribution may persist in a region because algal cover locally reduces the temperatures they experience, even when temperatures in surrounding unbuffered habitats exceed their physiological limits. Consequently, the continued presence of such species at a given latitude may not necessarily indicate that regional climatic conditions remain suitable, but rather that suitable microclimates are still being maintained by the persistence of habitat-forming algae. This buffering could therefore delay the apparent displacement of species while simultaneously increasing their dependence on the algae that provide these refugia. If warming eventually causes the loss or substantial reduction of these algae, the associated thermal refugia would disappear as well, potentially exposing multiple species to stressful conditions simultaneously [49]. Species responses to climate change may therefore be more abrupt than would be expected from a simple, linear relationship between regional temperature and species distributions, with a relatively gradual period of apparent persistence potentially followed by a rapid shift in community composition once the buffering habitat crosses its own ecological or thermal limits. Accounting for these dependencies and potential threshold responses will therefore be important when predicting future range shifts and community reorganization under climate change.

Another knowledge gap in the study of microclimate in coastal areas is that several local factors are rarely considered simultaneously, limiting assessment of their relative contributions. The cooling effect of cold microhabitats has been demonstrated in the southwest peninsula of England at mid-shore level [50] and for temperatures experienced by limpets of the genus *Patella* along the Iberian Peninsula [51]. Thermal buffering by *A. nodosum* canopies has been described in populations in the USA, but only up to 6 °C [52], whereas here we recorded differences of up to 20 °C. Our study shows the importance of both algal cover and solar exposure for the *in situ* temperatures experienced by organisms in the intertidal. Both factors were associated with thermal buffering of up to 20 °C especially relevant during high temperature peaks. Interestingly, some shore-specific variation occurred. While algal buffering generally resulted in cooling, at two shores in central Portugal the effect was reversed during summer in cold microhabitats, with higher temperatures recorded beneath the algae. This might be related to wind effects [53]. On a north-oriented, shaded surface (cold microhabitat), temperatures might remain low even during summer, especially during early morning after the tide has just receded. Evaporative cooling associated with wind might further reduce bare-rock temperatures, while below-canopy sensors are protected from wind exposure. Hence, the effect of wind and humidity needs to be explored in future studies to better describe the microclimate provided by canopies and the contribution of other local factors.

## Conclusions

Local-scale habitat complexity, particularly macroalgal cover, creates climate refugia that can modify the effects of broad-scale latitudinal gradients in rocky shores. While latitude had no detectable effect on intertidal temperatures across the six shores studied here, the low intertidal, the cold/shaded microhabitats and algal cover all provide significant cooling on rocky shores. The combined thermal buffering offered by biogenic and environmental factors (tidal level, solar exposure and algal cover) led to nearly 8.5 °C of variation in predicted experienced temperature. This thermal buffering might be key for the persistence of intertidal species [54] in the context of successive record-breaking temperatures in Portugal [55] especially considering that several species are at their southern distribution limits [10,24], and hence, likely very close to their upper physiological thermal limits [33]. Incorporating such microclimatic variation into predictions of future species distributions should improve our understanding of how rocky-shore communities will respond to continued warming.

## Acknowledgements

This work was supported by Fundação para Ciência e Tecnologia (FCT) through the projects BIOINTERACT (https://doi.org/10.54499/2022.02887.PTDC) and OceanLog (PTDC/BIA-BMA/4848/202). FPL and RdS received support through a CEECInd contracts (https://doi.org/10.54499/CEECIND/03185/2018/CP1546/CP1648/CT0004, https://doi.org/10.54499/2021.03268.CEECIND/CP1668/CT0001, respectively) and CMo through Foundation La Caixa (fellowship code: LCF/BQ/PR25/12110001). CMi, PDC and FA received support through FCT project UID/04292/2025 Centro de Ciências do Mar e do Ambiente (MARE), https://doi.org/10.54499/UID/04292/2025, and LA/P/0069/2020 granted to the Associate Laboratory ARNET., https://doi.org/10.54499/LA/P/0069/2020. The authors would like to acknowledge the support provided to PDC and CM by MARDIVE – Associação Ciência e Educação para a Conservação da Biodiversidade and FA by CMS.

## Supporting information

Fig. S1. Left: Temperature logger beneath *Fucus* sp. at Empa, central Portugal. Right: Removal of algal cover from a bare-rock treatment.

Fig. S2. Heatmap showing the classification of replicate pairs across all shores visits. Green: covered logger, score ≥4; yellow: covered logger, score=3; red: covered logger, score ≤2 or score ≥ 6, indicating no algal cover; grey: no data available. Bold font indicates the date of deployment. First column: shores ordered from north to south; second column: solar exposure (hot or cold).

Fig. S3. Boxplot showing the 25^th^ percentile, median, and the 75^th^ percentile of daily mean summer temperatures recorded on bare-rock in hot microhabitats, by shore and tidal level. Shores are ordered from north to south. Data on the left corresponds to the top intertidal and data on the right to the low intertidal. mol: Moledo; vcn: Viana do Castelo; emp: Empa; cbr: Cabo Raso; alt: Alteirinhos; ode: Odeceixe

Fig. S4. Scatter plots comparing the relationship between algal-cover buffering (temperature beneath algae minus temperature on bare rock) across the two tidal levels and two solar exposures (top-hot, top-cold, low-hot, and low-cold) at each shore. Shores are ordered from north to south, and summer data are shown. Red and blue dots represent daily mean low-tide temperatures during daytime and nighttime, respectively. The slope, confidence interval, R^2^ and p-value of each regression are reported in table S3.

Table S1. Selected replicate pairs and time periods used for the summer analysis.

Table S2 - Selected replicate pairs and time periods used for the winter analysis.

Table S3 - Linear regression analysis for algal-cover buffering against bare-rock temperatures by shore, tidal level, and solar exposure for the summer data. CI: confidence interval

## Notes

### Competing Interest Statement

The authors have declared no competing interest.

## References

1. Helmuth B, Mieszkowska N, Moore P, Hawkins SJ. Living on the Edge of Two Changing Worlds: Forecasting the Responses of Rocky Intertidal Ecosystems to Climate Change. Annu Rev Ecol Evol Syst. 2006;37: 373–404. doi:10.1146/annurev.ecolsys.37.091305.110149

2. Helmuth B, Broitman BR, Blanchette CA, Gilman S, Halpin P, Harley CDG, et al. Mosaic patterns of thermal stress in the rocky intertidal zone: implications for climate change. Ecol Monogr. 2006;76: 461–479. 10.1890/0012-9615(2006)076[0461:MPOTSI]2.0.CO;2

3. Smale DA, Wernberg T. Satellite-derived SST data as a proxy for water temperature in nearshore benthic ecology. Mar Ecol Prog Ser. 2009;387: 27–37. doi:10.3354/MEPS08132

4. Stobart B, Mayfield S, Mundy C, Hobday AJ, Hartog JR. Comparison of in situ and satellite sea surface-temperature data from South Australia and Tasmania: how reliable are satellite data as a proxy for coastal temperatures in temperate southern Australia? Mar Freshw Res. 2016;67: 612–625. doi:10.1071/MF14340

5. Meneghesso C, Seabra R, Broitman BR, Wethey DS, Burrows MT, Chan BKK, et al. Remotely-sensed L4 SST underestimates the thermal fingerprint of coastal upwelling. Remote Sens Environ. 2020;237: 111588.

6. Pereira L, Seabra R, da Silva R, Loureiro B, Lima FP. Autonomous, high-frequency and long-term monitoring of substrate temperature in rocky intertidal zones. Limnol Oceanogr Methods. 2026;24. doi:10.1002/lom3.10730

7. Kocsis ÁT, Reddin CJ, Kiessling W. The stability of coastal benthic biogeography over the last 10 million years. Glob. Ecol. Biogeogr. 2018;27: 1106–1120. doi:10.1111/GEB.12771

8. Moore NA, Morales-Castilla I, Hargreaves AL, Olalla-Tárraga MÁ, Villalobos F, Calosi P, et al. Temperate species underfill their tropical thermal potentials on land. Nat. Ecol. Evol. 2023 7:12. 2023;7: 1993–2003. doi:10.1038/s41559-023-02239-x

9. Boaventura D, Ré P, Cancela da Fonseca L, Hawkins SJ. Intertidal Rocky Shore Communities of the Continental Portuguese Coast: Analysis of Distribution Patterns. Mar. Ecol. 2002;23: 69–90. doi:10.1046/j.1439-0485.2002.02758.x

10. Monteiro C, Pereira J, Seabra R, Lima FP. Fine-scale survey of intertidal macroalgae reveals recent changes in a cold-water biogeographic stronghold. Front Mar Sci. 2022;9: 1–15. doi:10.3389/fmars.2022.880074

11. Lima FP, Ribeiro PA, Queiroz N, Hawkins SJ, Santos AM. Do distributional shifts of northern and southern species of algae match the warming pattern? Glob Chang Biol. 2007;13: 2592–2604. doi:10.1111/j.1365-2486.2007.01451.x

12. Lourenço CR, Zardi GI, McQuaid CD, Serrao EA, Pearson GA, Jacinto R, et al. Upwelling areas as climate change refugia for the distribution and genetic diversity of a marine macroalga. J Biogeogr. 2016;43: 1595–1607. 10.1111/jbi.12744

13. Monteiro C, Zardi GI, McQuaid CD, Serrão EA, Pearson GA, Nicastro KR. Canopy microclimate modification in central and marginal populations of a marine macroalga. Mar. Bio. 2017;49: 415–424. doi:10.1007/s12526-017-0824-y

14. Sousa MC, Ribeiro A, Des M, Gomez-Gesteira M, deCastro M, Dias JM. NW Iberian Peninsula coastal upwelling future weakening: Competition between wind intensification and surface heating. Sci. Total Environ. 2020;703: 134808. 10.1016/j.scitotenv.2019.134808

15. Wang D, Gouhier TC, Menge BA, Ganguly AR. Intensification and spatial homogenization of coastal upwelling under climate change. Nature 2015 518:7539. 2015;518: 390–394. doi:10.1038/nature14235

16. Nicastro KR, Zardi GI, Teixeira S, Neiva J, Serrão EA, Pearson GA. Shift happens: trailing edge contraction associated with recent warming trends threatens a distinct genetic lineage in the marine macroalga *Fucus vesiculosus*. BMC Biol. 2013;11: 6. doi:10.1186/1741-7007-11-6

17. Casado-Amezúa P, Araújo R, Bárbara I, Bermejo R, Borja, Díez I, et al. Distributional shifts of canopy-forming seaweeds from the Atlantic coast of Southern Europe. Biodivers Conserv. 2019;28: 1151–1172. doi:10.1007/s10531-019-01716-9

18. Herrero JJ, Simes DC, Abecasis R, Relvas P, Garel E, Ventura Martins P, et al. Monitoring invasive macroalgae in southern Portugal: drivers and citizen science contribution. Front Environ Sci. 2023;11: 1324600. doi:10.3389/FENVS.2023.1324600

19. Humet M, Pereira J, Nunes J, Nieto-Vilela R, Sinde-Mano A, Lima FP, et al. Proliferation of *Undaria pinnatifida* along the Atlantic coast of the Iberian Peninsula. Bot. Mar. 2025;68: 319–328. doi:10.1515/bot-2025-0013

20. Rindi L, Pedicini L, Bertocci I, Benedetti-Cecchi L, Smale D, Tuya F, et al. Variation in Environmental Conditions and Biodiversity Between Contrasting Intertidal Habitats Across Multiple Spatial Scales on European Rocky Shores. Glob. Ecol. Biogeogr. 2026;35: e70211. doi:10.1111/geb.70211

21. Smale DA, Teagle H, Hawkins SJ, Jenkins HL, Frontier N, Wilding C, et al. Climate-driven substitution of foundation species causes breakdown of a facilitation cascade with potential implications for higher trophic levels. J. Ecol. 2022;110: 2132–2144. doi:10.1111/1365-2745.13936

22. Bulleri F. Facilitation research in marine systems: state of the art, emerging patterns and insights for future developments. J. Ecol. 2009;97: 1121–1130. doi:10.1111/J.1365-2745.2009.01567.X

23. Coleman MA, Wernberg T. Forgotten underwater forests: The key role of fucoids on Australian temperate reefs. Ecol Evol. 2017;7: 8406–8418. 10.1002/ece3.3279

24. Pereira, Arenas F, Seabra R, da Silva R, Monteiro C, Pereira J, et al. Simulated intertidal heat stress on the brown seaweed *Ascophyllum nodosum* demonstrates differential population sensitivity to future climate. J. Ecol. 2025;113: 1504–1520. doi:10.1111/1365-2745.70037

25. Smale DA, Burrows MT, Evans AJ, King N, Sayer MDJ, Yunnie ALE, et al. Linking environmental variables with regional-scale variability in ecological structure and standing stock of carbon within UK kelp forests. Mar Ecol Prog Ser. 2016;542: 79–95. doi:10.3354/MEPS11544

26. Stelling-Wood TP, Poore AGB, Gribben PE. Shifts in biomass and structure of habitat-formers across a latitudinal gradient. Ecol Evol. 2021;11: 8831–8842. doi:10.1002/ece3.7714

27. IPMA. 2026. In: https://www.ipma.pt/pt/publicacoes/boletins.jsp?cmbDep=cli&cmbTema=pcl&idDep=cli&idTema=pcl&curAno=-1.

28. Thieurmel B, Elmarhraoui A. suncalc: Compute Sun Position, Sunlight Phases, Moon Position and Lunar Phase. CRAN: Contributed Packages. 2017. doi:10.32614/CRAN.package.suncalc

29. Harrison XA, Donaldson L, Correa-Cano ME, Evans J, Fisher DN, Goodwin CED, et al. A brief introduction to mixed effects modelling and multi-model inference in ecology. PeerJ. 2018;2018: e4794. doi:10.7717/PEERJ.4794

30. Schaefer N, Vadillo-Gonzalez S, Bishop MJ. Habitat-forming species as microclimatic refuges for intertidal organisms-a review and meta-analysis. Mar Environ Res. 2026;220: 108156. doi:10.1016/J.MARENVRES.2026.108156

31. Huang Y, Stein G, Kolle O, Kübler K, Schulze ED, Dong H, et al. Enhanced stability of grassland soil temperature by plant diversity. Nat. Geosci. 2023 17:1. 2023;17: 44–50. doi:10.1038/s41561-023-01338-5

32. Gril E, Spicher F, Vanderpoorten A, Gallet-Moron E, Brasseur B, Le Roux V, et al. The affinity of vascular plants and bryophytes to forest microclimate buffering. J. Ecol. 2025;113: 22–38. doi:10.1111/1365-2745.14425

33. Jurgens LJ, Gaylord B. Physical effects of habitat-forming species override latitudinal trends in temperature. Ecol Lett. 2018;21: 190–196. doi:10.1111/ELE.12881

34. Fiúza AFG. Upwelling Patterns off Portugal. In: Suess E, Thiede J, editors. Coastal Upwelling Its Sediment Record: Part A: Responses of the Sedimentary Regime to Present Coastal Upwelling. Boston, MA: Springer US; 1983. pp. 85–98. doi:10.1007/978-1-4615-6651-9_5

35. Borer G, Monteiro C, Lima FP, Martins FMS. Performance of DNA Metabarcoding vs. Morphological Methods for Assessing Intertidal Turf and Foliose Algae Diversity. Mol Ecol Resour. 2025;25: e14115. doi:10.1111/1755-0998.14115

36. Dijkstra J, Litterer A, Mello K, O’Brien B, Rzhanov Y. Temperature, phenology, and turf macroalgae drive seascape change: Connections to mid-trophic level species. Ecosphere. 2019;10. doi:10.1002/ecs2.2923

37. Miller RJ, Reed DC, Brzezinski MA. Community structure and productivity of subtidal turf and foliose algal assemblages. Mar Ecol Prog Ser. 2009;388: 1–11. doi:10.3354/MEPS08131

38. Martins GM, Hipólito C, Parreira F, Prestes ACL, Dionísio MA, José JM, et al. Differences in the structure and functioning of two communities: Frondose and turf-forming macroalgal dominated habitats. Mar Environ Res. 2016;116: 71–77. doi:10.1016/J.MARENVRES.2016.03.004

39. Smale DA. Impacts of ocean warming on kelp forest ecosystems. New Phytol. 2020;225: 1447–1454.

40. Wernberg T, Russell BD, Thomsen MS, Gurgel CFD, Bradshaw CJA, Poloczanska ES, et al. Seaweed Communities in Retreat from Ocean Warming. Curr. Biol. 2011;21: 1828–1832. 10.1016/j.cub.2011.09.028

41. Wernberg T, Filbee-Dexter K, de Bettignies T, Leclerc JC, Davoult D, Lévêque L, et al. Smaller plants in warmer water could have implications for future Kelp forests. Sci Rep. 2025;15: 28616-. doi:10.1038/s41598-025-13950-z

42. Serisawa Y, Akino H, Matsuyama K, Ohno M, Tanaka J, Yokohama Y. Morphometric study of *Ecklonia cava* (Laminariales, Phaeophyta) sporophytes in two localities with different temperature conditions. Phycological Res. 2002;50: 193–199. doi:10.1046/j.1440-1835.2002.00273.x

43. Pereira L, Morrison L, Shukla PS, Critchley AT. A concise review of the brown macroalga *Ascophyllum nodosum* (Linnaeus) Le Jolis. J Appl Phycol. 2020;32: 3561–3584. doi:10.1007/s10811-020-02246-6

44. Lalegerie F, Stengel DB. Concise review of the macroalgal species *Pelvetia canaliculata* (Linnaeus) Decaisne & Thuret. J Appl Phycol. 2022 34:6. 2022;34: 2807–2825. doi:10.1007/S10811-022-02830-Y

45. Lima FP, Wethey DS. Three decades of high-resolution coastal sea surface temperatures reveal more than warming. Nat Commun. 2012;3: 1–13.

46. Baptista V, Silva PL, Relvas P, Teodósio MA, Leitão F. Sea surface temperature variability along the Portuguese coast since 1950. International Journal of Climatology. 2018;38: 1145–1160. doi:10.1002/joc.5231

47. Pereira SC, Carvalho D, Rocha A. Temperature and Precipitation Extremes over the Iberian Peninsula under Climate Change Scenarios: A Review. Climate 2021, Vol 9, Page 139. 2021;9: 139. doi:10.3390/CLI9090139

48. Morelli TL, Daly C, Dobrowski SZ, Dulen DM, Ebersole JL, Jackson ST, et al. Managing Climate Change Refugia for Climate Adaptation. PLOS One. 2016;11: e0159909. doi:10.1371/journal.pone.0159909

49. Harley CDG, Paine RT. Contingencies and compounded rare perturbations dictate sudden distributional shifts during periods of gradual climate change. PNAS. 2009;106: 11172–11176. doi:10.1073/pnas.0904946106

50. Amstutz A, Firth LB, Spicer JI, Hanley ME. Facing up to climate change: Community composition varies with aspect and surface temperature in the rocky intertidal. Mar Environ Res. 2021;172: 105482. doi:10.1016/J.MARENVRES.2021.105482

51. Seabra R, Wethey DS, Santos AM, Lima FP. Side matters: microhabitat influence on intertidal heat stress over a large geographical scale. J Exp Mar Biol Ecol. 2011;400: 200–208.

52. Leonard GH. Latitudinal variation in species interactions: A test in the New England rocky intertidal zone. Ecology. 2000;81: 1015–1030. 10.1890/0012-9658(2000)081[1015:LVISIA]2.0.CO;2

53. Helmuth B, Yamane L, Lalwani S, Matzelle A, Tockstein A, Gao N. Hidden signals of climate change in intertidal ecosystems: What (not) to expect when you are expecting. J Exp Mar Biol Ecol. 2011;400: 191–199. doi:10.1016/J.JEMBE.2011.02.004

54. Jurgens LJ, Ashlock LW, Gaylord B. Facilitation alters climate change risk on rocky shores. Ecology. 2022;103: e03596. 10.1002/ecy.3596

55. IPMA. 2026. In: https://www.ipma.pt/pt/oclima/ondascalor/.

